# Accounting for imperfect detection reveals role of host traits in structuring viral diversity of a wild bat community

**DOI:** 10.1101/2020.06.29.178798

**Authors:** Anna R. Sjodin, Simon J. Anthony, Michael R. Willig, Morgan W. Tingley

**Affiliations:** Ecology and Evolutionary Biology, University of Connecticut, 75 N. Eagleville Road, Unit 3043, Storrs, Connecticut, USA 06269; Biological Sciences, University of Idaho, 875 Perimeter Drive, MS 3051, Moscow, Idaho, USA 83844; Center for Infection and Immunity, Columbia University, 722 W. 168^th^ Street, 17^th^ Floor, New York, New York, USA 10032; Center for Environmental Sciences & Engineering and Institute of the Environment, University of Connecticut, Storrs, Connecticut, USA 06269; Ecology and Evolutionary Biology, University of California – Los Angeles, 621 Charles E Young Drive South, Los Angeles, California, USA 90095

**Keywords:** cPCR, Chiroptera, Herpesvirus, Occupancy Modeling, Virus Community

## Abstract

Understanding how multi-scale host heterogeneity affects viral community assembly can illuminate ecological drivers of infection and host-switching. Yet, such studies are hindered by imperfect viral detection. To address this issue, we used a community occupancy model – refashioned for the hierarchical nature of molecular-detection methods – to account for failed detection when examining how individual-level host traits affect herpesvirus richness in eight species of wild bats. We then used model predictions to examine the role of host sex and species identity on viral diversity at the levels of host individual, population, and community. Results demonstrate that cPCR and viral sequencing failed to perfectly detect viral presence. Nevertheless, model estimates correcting for imperfect detection show that reproductively active bats, especially reproductively active females, have significantly higher viral richness, and host sex and species identity interact to affect viral richness. Further, host sex significantly affects viral turnover across host populations, as females host more heterogeneous viral communities than do males. Results suggest models of viral ecology benefit from integration of multi-scale host factors, with implications for bat conservation and epidemiology. Furthermore, by accounting for imperfect detection in laboratory assays, we demonstrate how statistical models developed for other purposes hold promising possibilities for molecular and epidemiological applications.

## 1. Introduction

Emerging infectious viruses, such as SARS-CoV 2 or Zika, represent single pathogen species embedded within multi-host, multi-pathogen systems. Community ecology – at the scale of host communities, and also at the scale of viral communities – is increasingly recognized as a productive framework with which to dissect such complex dynamics, illuminating risk factors associated with pathogen transmission, host switching, and spillover from wildlife to humans, among others [1, 2, 3, 4, 5, 6]. However, viral community ecology is limited by two major quantitative obstacles. First, most viruses are rare [7, 8, 9, 10, 11]. This means that large numbers of host individuals must be sampled for population-level detection, and resulting datasets are highly zero-inflated. Second, many wildlife viruses remain unknown to science [12, 13].

Currently, two of the most common methods for characterizing viral communities and finding new viral taxa are unbiased high-throughput sequencing (HTS; e.g. metagenomics) and consensus polymerase chain reaction (cPCR; [10]). HTS is advantageous when the goal is to construct an entire viral genome or when so little is known about the target taxon that primers cannot be designed. However, HTS lacks the sensitivity of PCR methods. cPCR is a method that relies on the use of degenerate primers for the broad amplification of both known and unknown viruses [10, 12]. It targets genome regions that are highly conserved among related viruses (e.g. at the family or genus level) and can detect multiple coinfecting viruses, if present. The degenerate primers bind with different efficiency to different viruses within a targeted taxon, so viruses with lower homology in the primer-binding region will not be detected as efficiently as would viruses with high sequence homology. Additionally, degenerate primers are, by definition, less specific. This reduces the sensitivity of cPCR and increases the probability of false negatives when viral load in the sample is low. This is a particular problem in wildlife surveys, where many samples are taken from healthy or asymptomatic animals [4, 14, 15, 16, 17, 18], and absence of symptoms is often associated with low viral load [19, 20, 21]. As a result, both HTS and cPCR methods have a high possibility of failed detection of true infections (i.e. false negatives).

Viral ecological studies using HTS and cPCR methods have mostly failed to account for the issues of false negatives. In general, high rates of false negatives are acknowledged (e.g. [4, 22]) but are rarely considered a major problem preventing honest inference. This becomes particularly problematic for studies of viral diversity and community assembly when accurate measures of the number and identity of detected viruses are necessary to infer patterns and mechanisms of co-infecting viruses. As consequent viral OTUs are missed for each sampled individual, the missing data compound when individuals are aggregated at the population and community levels. A handful of recent studies have used occupancy modeling techniques, adapted from wildlife ecology (where sites are surveyed to detect species, rather than where individual animals are assayed to detect viral strains), to better account for imperfect detection in an epidemiological context [23, 24, 25, 26]. However, as best we are aware, no one has yet adapted these methods to the detection process when using HTS or cPCR for biodiversity discovery or characterization of viral community ecology [27], and it is not a common practice to statistically account for failed detection in such a way across molecular biology more generally.

Additionally, viral ecologists have struggled to synthesize community dynamics across host scales. For example, infection heterogeneity among individuals is a key element in infectious disease dynamics [28, 29, 30] that is still relatively unexplored in viral communities of wildlife. Pathogen aggregation in host individuals can affect pathogen interactions, transmission, and host fitness, scaling up to alter population and community dynamics of the host. Failure to account for such heterogeneity has potentially dangerous or expensive consequences for modelling outbreak probabilities and forecasting disease spread [31, 32, 33].

Bats are an important host to understand from the perspective of viral transmission, as they seem to harbor a high number of viruses capable of infecting humans [34, 35, 36]. Cave-roosting bats in Puerto Rico are abundant and relatively easily captured, making them a practical system for addressing viral dynamics from an ecological perspective. Additionally, much is known about their behavior, biology, and life history that can guide contextual understanding of viral community dynamics [37]. For example, most cave-roosting species in Puerto Rico show sex-related spatial segregation during reproduction [37]. Females often use a centralized maternity colony within the cave, where they frequently and repeatedly come into contact with the same individuals. Males do not use this space. By continually re-exposing themselves to the same individual conspecifics, females likely reduce the variety of viruses to which they are exposed. Thus, host sex likely effects pathogen diversity. Indeed, the role of host sex has shown female-biased effects on ectoparasite and macro-parasite infection in Puerto Rican bats [38, 39]. Furthermore, many Puerto Rican caves are well-studied in terms of bat diversity and abundance. Two adjacent caves – Culebrones and Larva – have been studied extensively [37, 38, 39, 40, 41, 42, 43, 44, 45]. Species composition, as well as population sizes, diets, and reproductive phenology are all well understood for the bats roosting in these caves.

The goal of this study was to examine the role of host heterogeneity on viral diversity, while accounting for rarity and failed detection. To address these objectives, we collected wild bats from Culebrones and Larva Caves in Puerto Rico, tested bats for herpesvirus infection, and recorded individual-level host traits (mass, forearm length, ectoparasite intensity, sex, and reproductive status). To examine viral diversity across the scales of individual and population, we quantified a (individual-level) and γ (population-level) viral diversity, from which we calculated β diversity (compositional turnover of viruses among individual hosts).

Our research aimed to test multiple hypotheses in our inferential framework. First, we hypothesized that larger host individuals would have higher viral richness, as larger hosts provide more habitat space for pathogens and have shown higher levels of infection in previous work [46]. Second, we hypothesized that bats with more ectoparasites would harbor more viruses, as high parasite load and failure to groom have been shown previously to indicate illness or lowered immunity of hosts [47, 48, 49]. Third, we hypothesized that females and reproductively active individuals would have higher viral richness, as host sex [38, 39] and reproductive status [46, 50] have demonstrated impacts on viral infection in a diversity of host systems. Fourth, we hypothesized that bats roosting in the larger cave would have higher viral richness, due to greater host diversity and abundance in the larger cave. Relating to the community composition and diversity of infections, fifth, we hypothesized that β diversity of viruses would be greater in male than in female bats, as females limit their habitat range and use maternity colonies extensively during reproductive season, homogenizing their viral communities. Driven by relationships at the a and β levels, sixth, we hypothesized that γ diversity would be higher in female bats than male bats. Seventh, we hypothesized that γ diversity would be significantly different among host species, with species representing larger host populations having higher γ diversity than those roosting in smaller numbers. Finally, driven by strong relationships at the a and γ levels, we expected β diversity to differ among host species.

## 2. Materials and Methods

### (a) Sample Collection

Field work was conducted at Mata de Plátano Nature Reserve in Arecibo, Puerto Rico (18° 24.868’ N, 66° 43.531’ W). Mata de Plátano Nature Reserve (operated by InterAmerican University, Bayamon, Puerto Rico) is in the north-central, karstic region of Puerto Rico, an area dominated by caves suitable for roosting bats. Samples were collected from apparently healthy bats in Culebrones and Larva Caves for a total of 35 nights between June and August of 2017. Details about the cave habitats can be found in the Supplemental Material.

A harp trap, mist nets, and hand nets were used to capture 1,086 bats representing eight species: *Pteronotus quadridens, P. portoricensis, Mormoops blainvillii, Eptesicus fuscus, Monophyllus redmani, Brachyphylla cavernarum, Artibeus jamaicensis*, and *Erophylla sezekorni*. Upon capture, each bat was placed into a cotton holding bag, and individuals were identified to species based on Gannon et al. [37]. A clean cotton-tipped swab was used to collect saliva from each bat’s mouth. Swabs were placed in individual cryovials containing viral transport medium, maintained at −80° C in a dry shipper, and sent to Columbia University’s Center for Infection and Immunity. From each captured bat, we recorded six traits: mass, forearm length, ectoparasite intensity, sex, and reproductive status. Detailed descriptions of trait measurements can be found in the Appendix. Additionally, wing punches were taken from each bat for a separate analyses. To prevent re-sampling of recaptured bats, all subsequently captured bats with a punch-sized hole in the wing were released. All methods were approved by the University of Connecticut Institutional Animal Care and Use Committee (IACUC, protocol A15-032).

### (b) Laboratory Analyses

For each sample, herpesvirus cPCR assay targeting a ~170 base pair region of the catalytic subunit of the DNA polymerase gene of the herpesvirus family [51] was run twice (Figure 1). For assay one, all cPCR gel products (1% agarose) of expected size (Figure 1, b1) were cloned into Strataclone PCR cloning vector (Figure 1, c1), and twelve colonies were sequenced to detect viral co-occurrence and confirm viral identity via cross-referencing against the GenBank nucleotide database (Figure 1, d1). No cut bands from assay one resulted in false positives, so for assay two, only those gel products of expected size that were *not* present in assay one (Figure 1, b2) were cloned (Figure 1, c2) and sequenced (Figure 1, d2). Consequently, there were multiple samples from assay two for which data exist only on herpesvirus detection (positive or negative), but not on the identity of the detected herpesviruses (Figure 1, e2). Operational taxonomic units (OTUs) were determined using percent nucleotide identity and affinity propagation [11, 52] and were used in place of viral species.

**Figure 1:**
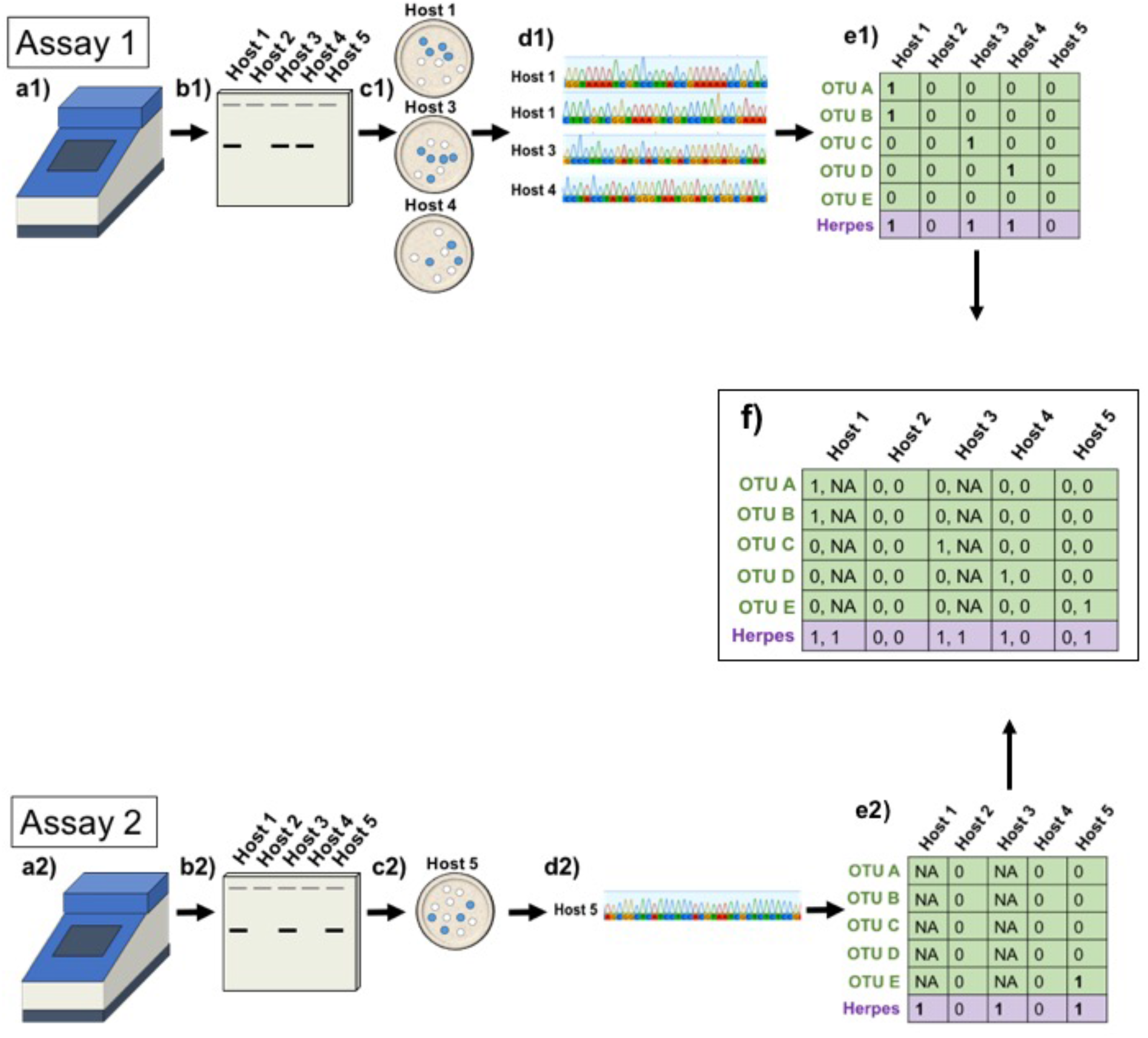
Depiction of molecular methods informing use of hierarchical community-level occupancy modeling. cPCR assays were run twice (a1 and a2). During assay 1, all gel bands were cut (b1), cloned (c1), and sequenced (d1). This resulted in OTU-specific presence-absence matrices that informed the response variable *y_i,j,k_*, presence of OTU *i* in individual *j* during assay *k*, for assay one (e1, green shading). These OTU-specific infections were then condensed to general herpesvirus (i.e. non-OTU-specific) presence-absence to inform the response variable *Y_j,k_*, presence of herpesvirus in individual *j* during assay *k* (e1, purple shading). For assay 2, only those gel bands that did not exist in b1 (e.g. Host 5, b2) were cut, cloned (c2) and sequenced (d2). This informed *y_i,j,k_*(e2, green shading) and *Y_j,k_* (e2 purple shading) for assay 2. Bands that existed in both assays (e.g. Hosts 1 and 3) were not cut, cloned, or sequenced during assay 2, and therefore OTU identity was not known. Consequently, for assay 2, these data only informed *Y_j,k_* (e2 purple shading). Both response variables for both assays (f) were used in the modeling framework.

Each sample was tested for PCR inhibition using TaqMan Exogenous Internal Positive Control Reagents (Applied Biosystems) with quantitative PCR as described in the manufacturer’s instruction manual. Amount of inhibition was determined using the standardized difference of sample quantitation cycle (Cq) from the mean Cq of two negative controls.

### (c) Modeling Framework

To explore the effects of individual-level host traits on herpesvirus infection, we fit a Bayesian community-level occupancy model to OTU-host occurrence data [53, 54]. This framework explicitly differentiates between the underlying state process determining the presence or absence of each OTU and the observation process determining the detection or non-detection of each OTU, conditional on its true presence. Although occupancy models have been employed in the field of animal ecology to study the occurrence of vertebrates across a landscape, the modeling framework has rarely yet been applied in molecular settings to account for imperfect detection during indirect sampling, such as with eDNA [55, 56] or in studies of pathogen ecology [23, 24, 25].

By modeling OTUs in a community context, we improved accuracy of estimates of incidence for rare OTUs by considering parameter estimates as all coming from a similar stochastic process, a Bayesian process known as “borrowing strength” [57, 58]. As a trade-off, the model does not robustly inform which host traits shape occupancy of individual OTUs, but provides strong inference on host trait relationships to the aggregate community of OTUs.

In the model, response variable *y_i,j,k_*, is a binomial random variable denoting whether herpesvirus OTU *i* was detected (*y_i,j,k_* = 1) or not (*y_i,j,k_* = 0) in host individual *j* during assay *k* (green values in Figure 1e and f). A second response variable, *Y_j,k_*, is a binomial random variable denoting whether any herpesvirus, regardless of OTU identity, was detected (*Y_j,k_*, = 1) or not (*Y_j,k_*, = 0) in host individual *j* during assay *k* (purple values in Figure 1 e and f). The two response variables, *y* and *Y*, are hierarchically related, yet independently observed via previously described methods: cPCR identifies the presence or non-detection of any herpesvirus (*Y*), and subsequent sequencing and bioinformatics identifies the presence or non-detection of particular OTUs (*y*).

The model simultaneously estimates the probability of infection (*ψ_i,j_*) and the probability of detection (*p_i,j,k_*) with a mixture model in the form *y_i,j,k_* ~ Bernoulli (*p_i,j,k_* * *z_ij_*), where *p_i,j,k_* is the probability of detecting OTU *i* in host individual *j* during assay *k*, and *z_ij_* is a latent variable representing true infection of OTU *i* in host individual *j*, deriving from the equation *z_ij_* ~ Bernoulli (*ψ_ij_*,). A second latent variable (*Z_j_*) represents true herpesvirus infection, regardless of OTU identity, in host individual *j*. The model’s hierarchical structure allows the two response variables (*y_i,j,k_* and *Y_jk_*), as well as the two latent variables (*z_ij_*, and *Z_j_*), to inform one another, as *Z_j_* = max (*z_ij_*) and *Y_j,k_*, ~ Bernoulli (*p_assay_* * *Z_j_*), where *p_assay_* represents a constant probability that the cPCR assay will detect a herpesvirus given the presence of at least one OTU. Together, the model assumes that an empirical detection (*y_ijk_* = 1, *Y_jk_* = 1) represents true viral infection, but that an empirical non-detection (*y_i,j,k_* = 0, *Y_j,k_* = 0) could indicate either a true absence or a true infection with failed detection.

To structure the model, sample inhibition and OTU subfamily were included as fixed effects influencing OTU-specific detection probability (*p_i,j,k_*) Host forearm length, cave identity, ectoparasite intensity, sex, reproductive status, and the interaction between sex and reproductive status were included as fixed effects for infection probability (*ψ_i,j_*). Host species was included as a random intercept for both detection and infection probabilities.

As is typical for community occupancy models, every intercept and slope parameter for a given herpesvirus was assumed to be drawn from a hierarchical Normal distribution that describes the distribution of slopes for all herpesvirus species available for sampling [59]. In this way, primary inference on the community is made by the examination of the hyper-parameters that provide a mean community-wide effect for any given covariate. As preliminary explorations of the dataset indicated that some herpesviruses were unusually common or unusually rare, instead of using a hierarchical Normal to describe the random distribution of occupancy intercepts, we used a fat-tailed distribution, the *t*-distribution. Full model code is provided online as Supplemental Material.

We executed the model in JAGS [60] using the R2jags package [61] with vague priors (e.g., normal with μ = 0, τ = 0.1, gamma with shape and rate = 0.01). We ran three chains of 15,000 iterations thinned by 50 with a burn-in of 25,000, yielding a posterior sample of 900 iterations across all chains. We checked convergence visually with traceplots and made parameter inference based on 95% Bayesian credible intervals. Posterior predictive checks were conducted for individual OTUs by calculating Bayesian p-values [62] and 90% Bayesian credible intervals for mean detected prevalence of each OTU. A lack of fit for a particular OTU would be indicated by a 90% credible interval that did not overlap the empirically observed prevalence. All analyses were done using R version 3.4.3 [63].

### (d) Alpha, Beta, and Gamma Partitions

To examine viral diversity across the levels of host individual and population, we compared α, β, and γ diversity among host species and between host sexes using 900 host-by-OTU matrices drawn from the community occupancy model’s posterior distribution of the true underlying infection state, *z_i,j_*. α diversity was calculated as the average viral richness per individual bat across 1) all species for each sex, 2) both sexes for each species, and 3) each species-by-sex combination (male *A. jamaicensis*, female *A. jamaicensis*, male *P. quadridens*, etc.). γ diversity was defined as the total viral richness for 1-3. β diversity was calculated using both the multiplicative (γ = α * β; [64]) and additive (γ = α + β; [65]) models.

For α diversity, the general statistical design is that of a two-factor analysis of variance (ANOVA) with replication, with individual bats representing replicates. To test the significance of results, we calculated F-statistics for differences in a diversity associated with 1-3. We then compared empirical F-statistics for each of the 900 posterior draws to three null distributions (H0: no difference in α diversity between host sexes or among host species). Each null distribution was generated using a different assumptions regarding the relationship between relative abundances of each sex-by-species group, as compared to the true proportion of individuals in that group. Details describing generation of the null distribution are available in the Appendix.

For β and γ diversity, the general statistical design is that of a two-factor ANOVA without replication, with treatment groups representing replicates. As such, values were only compared between sexes and among species (1 and 2). We could not assess their interaction.

Simulation methods were identical to those for the a-level analyses (Appendix). All analyses were done using R version 3.4.3 [63]. R code used for the simulation methods and calculation of F-statistics is available at github.com/ARSjodin/AlphaBetaGamma.

### 3. Results

#### (a) Model Fit

Screening results and OTU delineations are reported previously [11]. Posterior predictive checks on each individual OTU indicate an overall adequate degree of model fit, with no 90% credible intervals falling outside of the observed prevalence for an OTU (Supplemental Table 2). Observed prevalence was more likely to be underpredicted than overpredicted, indicating that the model may be more pessimistic about detection probability than observed in the real dataset.

#### (b) Imperfect detection of OTUs

Model fits indicated OTUs were imperfectly detected at all stages of laboratory analysis. The cPCR assay had a 0.778 probability (*p_assay_*, 95CrI = 0.744–0.807) of detecting a herpesvirus given the presence of at least one OTU (Table 1). Given two cPCR assays, this suggests a 0.049 probability (95CrI = 0.037–0.065) of incorrectly assuming a host is not infected, were detection assumed to be perfect. However, even if the cPCR assay suggests a positive infection, not every OTU will be detected via sequence amplification. On average across all OTUs, the probability of detecting a single OTU via sequencing was 0.699 (inverse logit-transformed *μ*_*a*0_, 95CrI = 0.637–0.762). There was no support for a role of either sample inhibition or viral subfamily in affecting viral detection rates (Table 1). Combined across both stages of viral detection, there was both a non-zero probability that cPCR assays would fail to detect a viral infection, and that sequence amplification would then fail to identify the full suite of OTUs present. Consequently, the fitted model estimated on average that 1.75 hosts (95CrI = 0 – 5) likely held infections of at least one OTU that were not detected by cPCR, and that the mean number of OTUs missed per host was 0.036 (95CrI = 0.017–0.063). Across the tested host community (n = 41), this combined to an expected undetected total of 12.2 OTU infections (95CrI = 5.5–21.0).

**Table 1.** Estimates of hyper-parameter means and their upper and lower 95% credible intervals for the community occupancy model of herpesvirus occurrence in bat hosts. Hyper-parameters represent the average community-wide effect across all viral OTUs, and estimates are presented on the logit scale. Estimates for slope parameters with credible intervals that do not overlap zero are presented in bold.

| Response | Hyper-parameter | Estimate | Lower CrI | Upper CrI |
| --- | --- | --- | --- | --- |
| Detectability | Intercept | 0.840 | 0.549 | 1.168 |
| - | Inhibition | -0.065 | -0.343 | 0.210 |
| - | Sub-family | 0.067 | -0.560 | 0.715 |
| Occupancy | Intercept | -8.702 | -10.725 | -6.720 |
| - | Cave | -0.208 | -1.911 | 1.112 |
| - | Sex | 0.302 | -0.058 | 0.643 |
| - | **Reproductive Status** | 1.394 | 0.384 | 2.354 |
| - | **Sex \* Reproductive Status** | -1.166 | -1.912 | -0.510 |
| - | Forearm Length | -0.041 | -0.130 | 0.047 |
| - | Ectoparasite Intensity | -0.197 | -0.489 | 0.052 |

#### (c) Role of Host Traits

In general, herpesvirus occupancy was low in Puerto Rican bats, but the model uncovered robust support for the importance of individual-level traits as drivers of viral richness (Table 1). Herpesvirus richness was higher in reproductively active bats compared to non-reproductive bats. Specifically, reproductively active female bats had higher viral richness as compared to both non-reproductive bats and to reproductively-active male bats. No other host traits significantly affected the probability of viral occurrence.

Using model estimates of true occurrence correcting for imperfect detection, bats showed differing degrees of herpesvirus diversity across community scales. a diversity was significantly different between the sexes, but this effect differed by species, with only some bat species showing sex-specific differences in a diversity (Figure 2, Appendix Table 3). Despite strong differences in a diversity by species and sex, this did not scale up to g-level differences (Figure 2, Appendix Table 3). Taken together, this means that although certain bat species (and sexes within those species) host more OTUs than do other species, the identity of the OTUs is quite similar across individuals of a species, indicating low OTU turnover at the population level. Indeed, β diversity did not differ significantly across species (Figure 2, Appendix Table 3). However, the multiplicative form of β diversity (i.e. γ = α * β) was significantly different between sexes, with higher β diversity, and therefore higher turnover of OTUs, in females than males (Figure 2, Appendix Table 3). Because no γ-level difference existed between bat sexes, and the sex differences at the a level depended on species, the differences in β diversity between sexes is likely driven by α-level effects of certain species.

**Figure 2:**
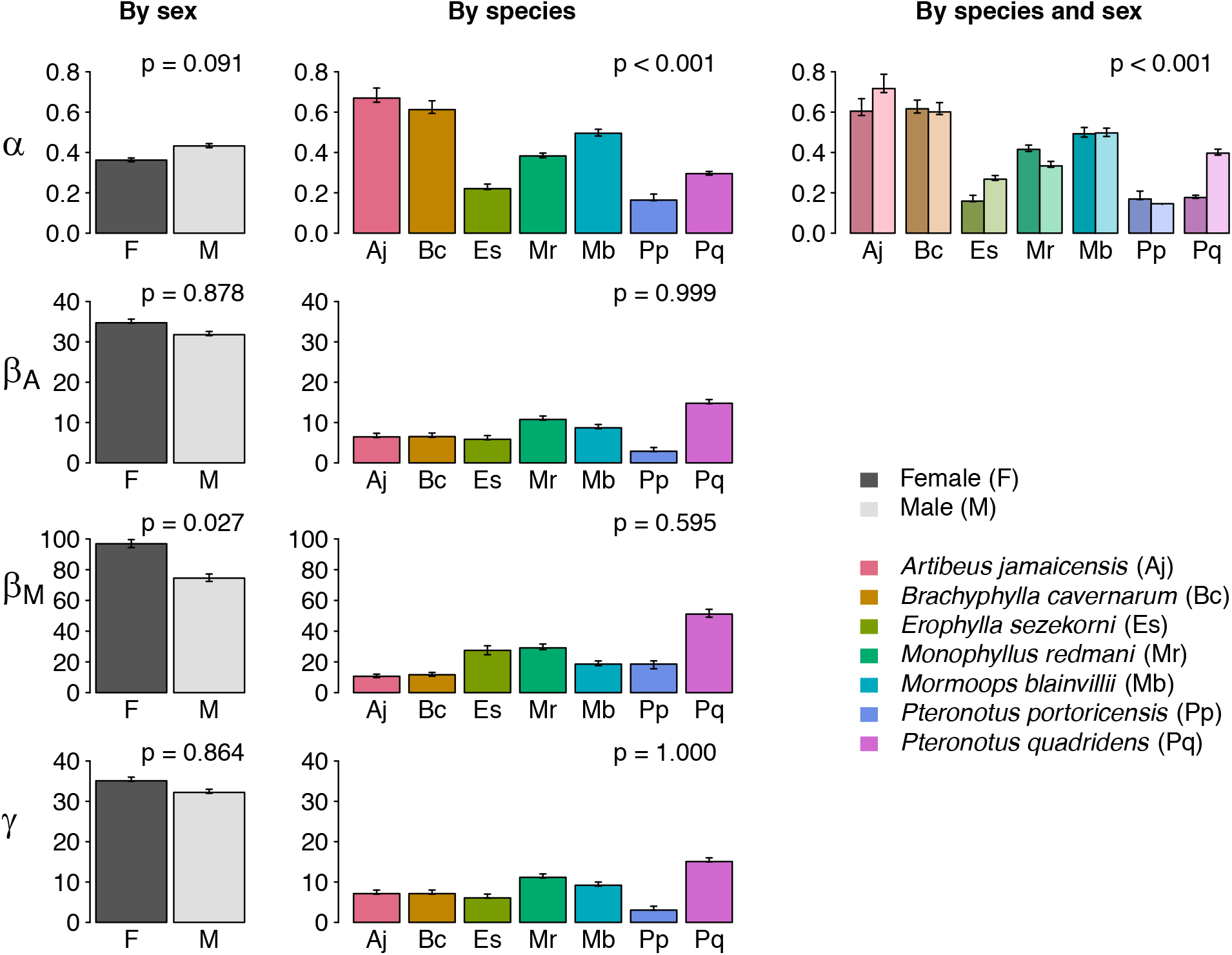
α, β, and γ diversity by sex (black and grey shading), species (color), and species-by-sex groups. Rows represent diversity metrics (β_M_ signifies the multiplicative model, and β_A_ signifies the additive model). Error bars indicate posterior uncertainty in metrics calculated using 900 z-matrices. P-values represent significance when comparing empirical metrics to a null distribution using the two-step null generation method (see Appendix). P-values associated with comparisons using two additional null distributions, which do not show substantive differences, are included in Appendix Table 3.

## 4. Discussion

Scientists and public health officials have called for a shift from post-emergence, reactive viral studies to predictive, preventative approaches [66, 67]. However, the integration of disease ecology and ecological forecasting is still in its infancy and is constrained in utility [67, 68, 69, 70, 71, 72, 73]. An ongoing hindrance to progress is data latency, or the lag time between data collection and data availability for analyses [72]. In the context of understanding viral shedding in wildlife, this includes the time spent capturing and sampling hosts in the field, transporting samples from the field to the laboratory, extracting genetic material and diagnosing infection, and subsequent sequencing and bioinformatics. By linking increased viral shedding to more easily and quickly attainable host attributes such as reproduction and the interaction between sex and reproductive state, our study suggests that the use of host traits may serve as a proxy for transmission risk, providing a potential solution to the data latency problem. For example, in many bat species, including those tested here [37], reproductive status is a trait that is predictable based on season, making it a more useful proxy in forecasting [74]. Assuming this relationship holds for other virus-host systems, when scientists make forecasts about the risk posed by bats to human health, reproductive status could potentially be added to models and updated temporally without extensive, ongoing molecular sampling. Ongoing investigations the relationship between viruses and host reproduction would help illuminate the utility of this proxy.

Although all living organisms contain a multitude of viruses [34, 75, 76], targeted testing of bats, and the resulting diversity of discovered viruses, have created conflicting perceptions about the threat or value of bats to humans. Dramatized communication of bat surveillance has exacerbated the perceived negative reputation of bats, causing viral discovery efforts to become a conservation issue [77]. Unfortunately, efforts to intentionally kill wild bats for protection of human health have threatened populations worldwide [78, 79, 80]. However, the effect of reproduction on viral shedding, as presented here, suggests non-contradictory goals of bat conservation and public health. Because bats, especially female bats, shed more viruses during reproduction, the risk of viral transmission from bats to humans is higher during this time. Additionally, reproductive rate is an intrinsic factor that informs bat conservation [81] and correlates with extinction rate [82]. Disturbances caused by humans during reproduction can increase mortality rates of offspring [84], which can then cause population declines and threaten population persistence. Therefore, minimizing human disturbance to bats during reproduction, especially to maternity colonies, can curtail the risk of viral transmissions to humans and human-induced declines of bat populations.

At the level of individual hosts (α), host sex and species interacted to significantly affect viral diversity (Table 2). This suggests that the difference in a diversity among the host species is different for males and females of each species. This could be a result of an antagonistic relationship between host species and sex (e.g. males of species one have *higher* viral diversity than females of species one, but males of species two have *lower* viral diversity than females of species two). Alternatively, this could be a result of differing magnitude of viral diversity among host species and sex (e.g. males of species one have *much* higher viral diversity than females of species one, but males of species two have only *slightly* higher viral diversity than females of species two). Values of α diversity (Figure 2, Appendix Table 3) suggest that the differences in viral richness between sexes generally differ in magnitude among host species, and the impact of sex is not antagonistic among species.

Host sex, alone, had a significant effect on β diversity (using the multiplicative model), with female bats hosting more compositionally heterogeneous infracommunities (i.e. higher β diversity; Table 2, Figure 2). This was contrary to our hypothesis and shows that, despite limiting range size and repeated exposure to the same individual conspecifics, the viral metacommunity of female bats does not become homogenized compositionally. Instead, compositional heterogeneity could be a result of strong physiological differences between immune-competence of the sexes, especially during reproduction. Immune suppression of reproductively active female bats may activate previously latent herpesviruses, resulting in viral infracommunities that comprise OTUs acquired at any point in time. However, this could not be tested in our study. If supported, it would suggest that viral composition in female bats may not be as temporally constrained as in males, resulting in a more compositionally heterogeneous metacommunity.

Notably, differences in β diversity related to host sex are shaped strongly by two host species, *P. quadridens* and *P. portoricensis* (Appendix Table 3). Unfortunately, little is known about behavioral or physiological differences between the sexes of each of these species, that may contribute to viral sharing in a manner that is different from that of other species. Investigations of the mechanistic relationship between host physiology and viral infection is a fruitful area for further research. Additionally, this difference in viral β diversity between male and female *P. quadridens* is due primarily to differences in α diversity between the sexes, as γ diversity of male and female *P. quadridens* are indistinguishable (Appendix Table 3). For *P. portoricensis*, the relationship is reversed. These results further emphasize the need to explicitly incorporate both individual- and population-level heterogeneity in analyses of viral ecology.

In terms of viral detection, neither sample inhibition nor OTU subfamily influenced overall viral detection rates (Table 1). The near-zero effect size of sample inhibition is important because it demonstrates that samples do not contain contamination or non-viral genetic material that prohibited detection of herpesviruses. Additionally, the near-zero effect size of OTU subfamily shows that the assay is not biased towards either the Beta- or Gammaherpesvirinae. Notably, no alphaherpesviruses were detected in the samples [11], so the results make no suggestions about the assay’s efficiency in detecting viruses belonging to this subfamily. Still, detection was imperfect, and reasons driving failed detection are yet to be determined. One unmeasured factor that may influence the probability of detection in this system is viral load, or the intensity of viral infection (i.e. viral “abundance” in each host). Indeed, infection intensity is a significant driver of detection probability in other disease systems [24, 25]. In the context of the present study, however, quantifying viral intensity in each sample would require development of separate quantitative PCR assays for each OTU and was beyond the scope of this work.

Failed detection in this system can occur at the stage of sample collection (i.e. the host is infected, but the virus is not in the sample) and at the stage of molecular analyses (i.e. the virus is in the sample, but the laboratory methods did not detect it). Reasons could be biological (e.g. low viral load associated with asymptomatic hosts) or methodological, if tests fail because of genetic variation in the primer-binding sites or some other mechanism, such as failure to capture low-level sequences in the cloning step. The methods used here only account for failed detection at the stage of molecular analyses – as only one sample was tested per host – thus failed detection at the stage of sample collection will be perceived here as a lack of occurrence. However, our modeling method could easily be expanded to accommodate the former, additional source of heterogeneity, by taking and testing multiple samples per host.

Our study illustrates how viral community ecology can be used to identify risk factors for infection while quantitatively accounting for imperfect detection, serving as a model for studies of viral communities moving forward. Community-level occupancy modeling can be applied to continued viral discovery efforts, molecular biology more broadly, and to applications of approaches or concepts from community ecology to virus systems. For the herpesvirus assay used in this study [51], the model indicated that two assay runs were able to detect approximately 95% of true infections. A third run would have resulted in an estimated 99% detection. Similar calculations, following [84], can be done for other assays to guide the design of molecular methods. Additionally, the development of a novel hierarchical model that uses a single assay run with specific OTU identification (Figure 1, e1), followed by assay runs that detect infection only, without OTU identification (Figure 1, e2), accounts for false absences without considerably increasing sequencing costs or time spent analyzing additional sequences. The further co-development of emerging quantitative models with molecular techniques stands to improve both the strength of findings as well as the cost-efficiency of assays.

## Acknowledgements

We thank Armando Rodríguez-Duran for logistical help in the field, as well as the field team, including Sarah Stankavich, Jose Rivera, Melanie Hodge, Brian Springall, Elspeth Pierce, Emily Stanford, Juan Contreras, Ismael Ribot, and Frank Sjodin. Isamara Navarrete-Macias and Eliza Liang trained ARS on laboratory methods, and Heather Wells provided lab and bioinformatics help. Eliza Grames provided invaluable coding assistance. We also thank Janine Caira, Sarah Knutie, and Mark Urban for constructive discussions that advanced the intellectual development of the work. A.R.S. was funded by the NSF Graduate Research Fellowship Program and a Jorgensen Fellowship from the University of Connecticut. Both A.R.S. and M.W.T. additionally received support from The Center for Biological Risk at the University of Connecticut. M.R.W. was funded by a grant from NSF (DEB-1546686 and DEB-1831952). Fieldwork was supported by Sigma Xi; the Royal Society of Tropical Medicine and Hygiene; and the University of Connecticut’s Department of Ecology and Evolutionary Biology, El Instituto via the Tinker Foundation, and the Office of the Vice President for Research. This project also benefited from support from the USAID PREDICT project.

## Author Contributions

A.R.S. conducted field and laboratory work. S.J.A. and A.R.S. designed laboratory methods. M.W.T. and A.R.S. designed modeling methods. M.R.W. and A.R.S. designed simulation methods. M.W.T. and A.R.S. conducted quantitative analyses and drafted the manuscript. All authors provided comments and gave final approval for publication.

## Description of cave habitats

The larger cave (Culebrones Cave) is a structurally complex hot cave, with temperatures reaching 40° C and relative humidity at 100%. It is home to an estimated 300,000 bats representing six species: *Pteronotus quadridens, P. portoricensis, Mormoops blainvillii, Monophyllus redmani, Erophylla sezekorni*, and *Brachyphylla cavernarum* [1]. The smaller cave (Larva Cave) is a cold cave that is at ambient temperature and less structurally complex compared to Culebrones Cave. It hosts fewer bats (30-200, ARS personal observation), representing two species, *Artibeus jamaicensis* and *Eptesicus fuscus*. No bat species roosting in the larger cave was found roosting in the smaller cave, and vice versa.

## Description of trait measurements

Bats were weighed to the nearest 0.5 grams using a Pesola scale. Using calipers, forearm length was measured to the nearest 0.01 millimeter from the distal tip of the radius to the distal tip of the olecranon process. Ectoparasite intensity was measured from visual inspection of the number of individual ecotparasites and represented as a categorical variable: low (< 5), medium (5–8), high (9–19), and extreme (>19). A categorical measure of intensity was used instead of abundance because best practices to quantify exact abundance require host anesthesia [2]. Voucher specimens were collected for each ectoparasite morpho-species and classified broadly as flies from the Hippoboscoidea superfamily, mites, and ticks, each of which had multiple morpho-species and variable host ranges (Appendix Table 1). Specimens were deposited in the University of Connecticut’s Biodiversity Research and Education Collections. For female bats, pregnancy was determined via gentle palpation of the abdomen, and lactation was determined by milk expression or visible mammary formation. Pregnant or lactating bats were considered to be reproductively active. For male bats, reproductive activity was determined by the presence of descended testes.

## Model code

The following represents statistical code written in the JAGS modeling language for fitting the multi-species occupancy model of viral community occurrence.

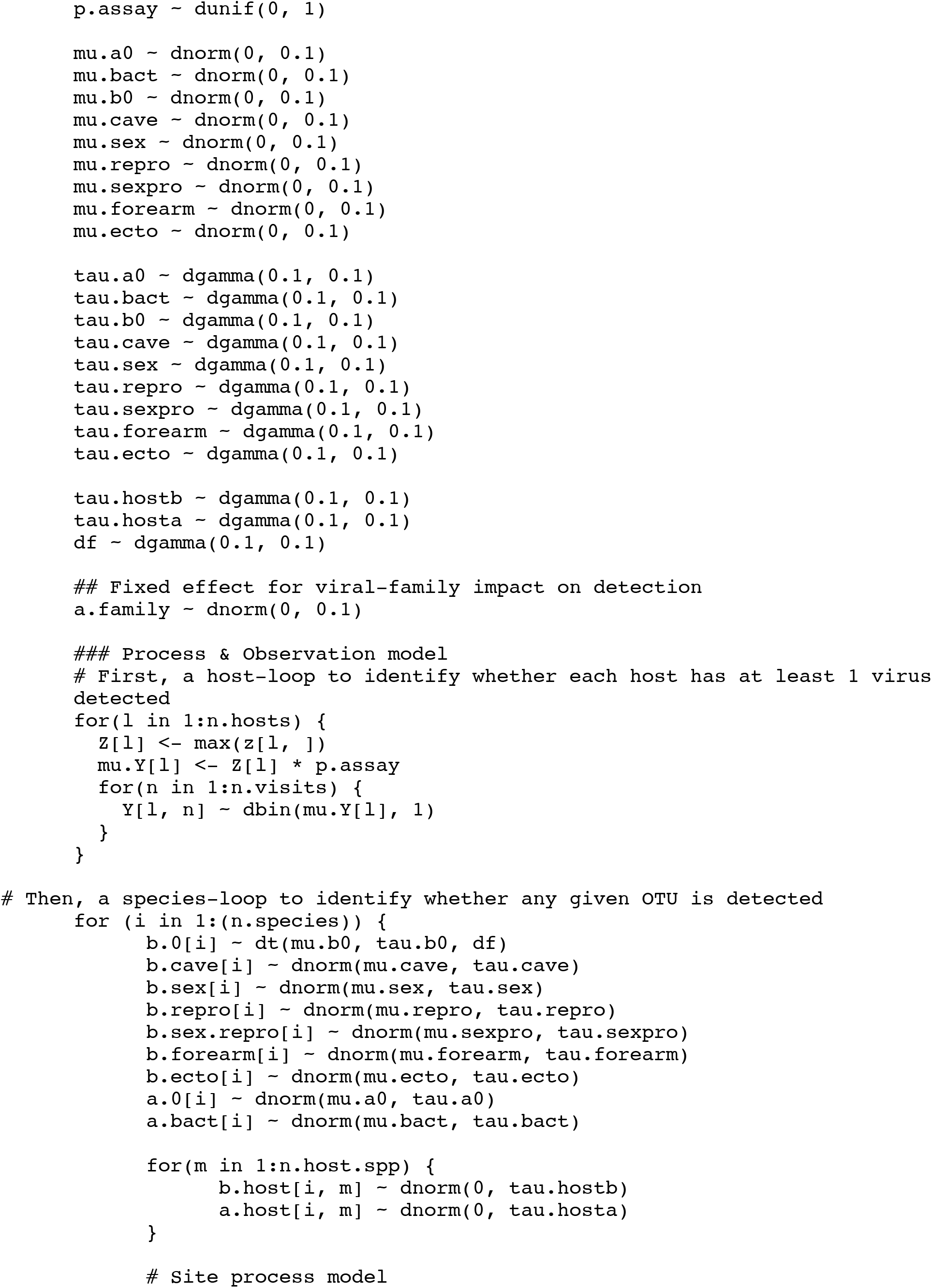
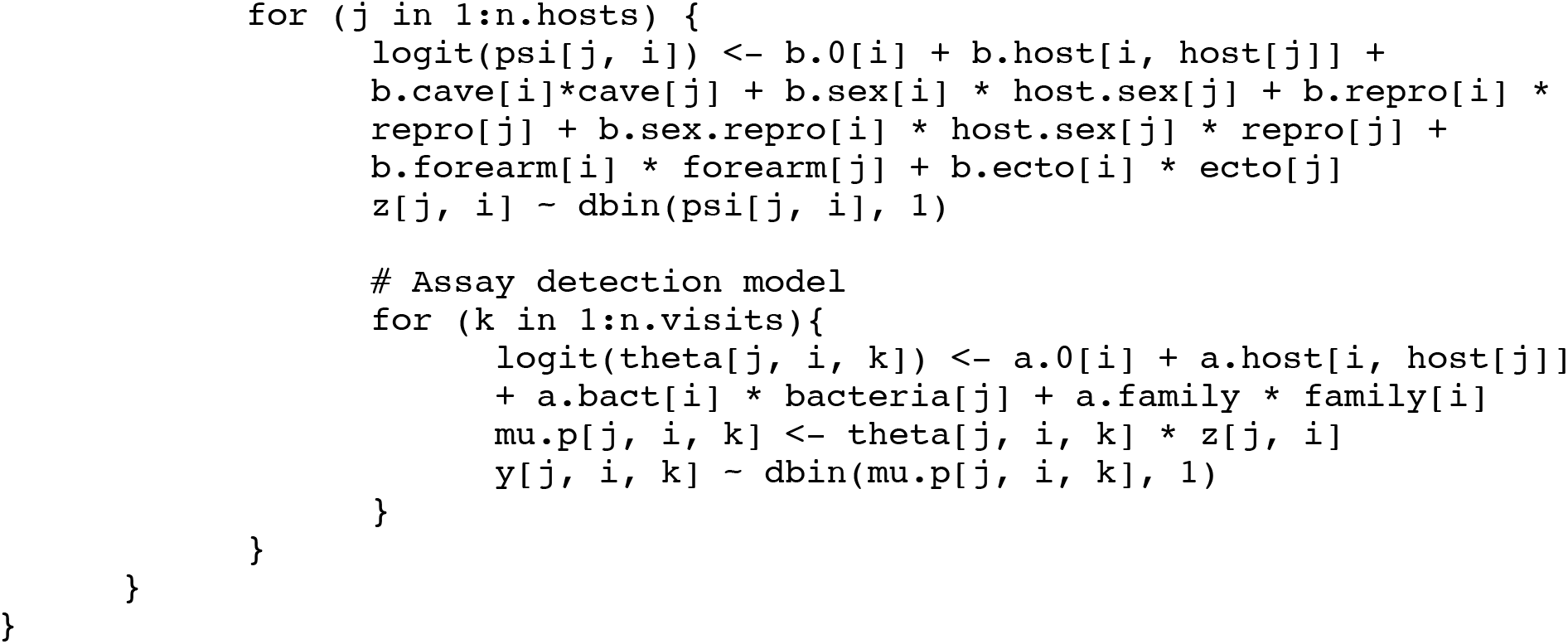

## Generation of null distribution

To create the null distribution, all empirical viral richness values from all sampled bats were placed into a pool (Appendix Figure 1). Richness values from individual bats were then selected at random, and without replacement, from the pool and randomly assigned a host treatment. Random draws continued until simulated treatment groups contained the same number of bats (N) as do empirical treatment groups.

This simulation method assumes that the empirical relative abundances of each host treatment group represent the true proportion of individuals of that treatment. We evaluated the robustness of results under this assumption by expanding the methodological approach in two ways. First, we repeated the above methods, but sampled from the regional pool *with* replacement. Second, we adopted a two-part simulation in which step one included random selection of a host species by sex treatment as a “donor”, such that each treatment combination had the same chance of providing hosts to the simulated data set. In step two, we randomly drew an infracommunity from the previously selected donor treatment and placed it into a randomly selected “recipient” treatment. The donor treatments were sampled with replacement, and the two-step simulation was repeated until the number of infracommunities in each recipient treatment equaled empirical numbers.

**Appendix Table 1:** Ectoparasite morpho-species and their associated hosts

| Morpho-Species | Host Species | Infection Site | Number of Infected Hosts |
| --- | --- | --- | --- |
| Hippoboscoidea Fly 1 | *Artibeus jamaicensis* | Fur | 20 |
| Hippoboscoidea Fly 2 | *Artibeus jamaicensis*, *Brachyphylla cavernarum*, *Eptesicus fuscus*, *Erophylla sezekorni*, *Monophyllus redmani*, *Mormoops blainvillii*, *Pteronotus portoricensis*, *P. quadridens* | Fur | 210 |
| Hippoboscoidea Fly 3 | *Monophyllus redmani*, *Pteronotus portoricensis*, *P. quadridens*, | Fur | 202 |
| Mite 1 | *Artibeus jamaicensis*, *Monophyllus redmani* | Ears | 51 |
| Mite 2 | *Pteronotus quadridens* | Wings | 133 |
| Mite 3 | *Artibeus jamaicensis*, *Brachyphylla cavernarum*, *Eptesicus fuscus*, *Erophylla sezekorni*, *Monophyllus redmani*, *Mormoops blainvillii*, *Pteronotus portoricensis* | Wings | 209 |
| Tick 1 | *Erophylla sezekorni*, *Monophyllus redmani* | Furred body | 31 |
| Tick 2 | *Pteronotus quadridens* | Membranes | 3 |

**Appendix Table 2:** Observed vs. predicted OTU prevalence, ranked from most prevalent to least prevalent OTU. Bayesian p-values provide estimates of model fit, with values < 0.05 indicating potential lack of fit. In general, the model showed good fit for most OTUs, with evidence of mild lack-of-fit (under-estimation of prevalence) for the rarest and most common OTUs.

| OTU | Observed Prevalence | Mean Predicted Prevalence | Bayesian p-value |
| --- | --- | --- | --- |
| 21 | 0.062 | 0.058 | 0.059 |
| 4 | 0.058 | 0.053 | 0.052 |
| 5 | 0.037 | 0.034 | 0.163 |
| 18 | 0.029 | 0.027 | 0.193 |
| 1 | 0.028 | 0.024 | 0.057 |
| 12 | 0.016 | 0.014 | 0.067 |
| 9 | 0.015 | 0.014 | 0.407 |
| 14 | 0.013 | 0.012 | 0.373 |
| 24 | 0.013 | 0.012 | 0.432 |
| 27 | 0.012 | 0.011 | 0.351 |
| 34 | 0.012 | 0.011 | 0.427 |
| 23 | 0.0083 | 0.0073 | 0.382 |
| 28 | 0.0074 | 0.0067 | 0.499 |
| 13 | 0.0064 | 0.0061 | 0.342 |
| 16 | 0.0064 | 0.0060 | 0.408 |
| 19 | 0.0055 | 0.0052 | 0.379 |
| 20 | 0.0055 | 0.0052 | 0.333 |
| 30 | 0.0055 | 0.0052 | 0.350 |
| 37 | 0.0055 | 0.0050 | 0.494 |
| 11 | 0.0055 | 0.0043 | 0.351 |
| 6 | 0.0037 | 0.0035 | 0.272 |
| 2 | 0.0028 | 0.0025 | 0.303 |
| 7 | 0.0028 | 0.0025 | 0.303 |
| 15 | 0.0028 | 0.0025 | 0.321 |
| 17 | 0.0028 | 0.0026 | 0.222 |
| 22 | 0.0028 | 0.0026 | 0.239 |
| 38 | 0.0028 | 0.0026 | 0.246 |
| 39 | 0.0028 | 0.0026 | 0.260 |
| 25 | 0.0018 | 0.0017 | 0.156 |
| 29 | 0.0018 | 0.0017 | 0.169 |
| 32 | 0.0018 | 0.0017 | 0.162 |
| 40 | 0.0018 | 0.0017 | 0.207 |
| 3 | 0.00092 | 0.00086 | 0.110 |
| 8 | 0.00092 | 0.00083 | 0.132 |
| 10 | 0.00092 | 0.00086 | 0.098 |
| 26 | 0.00092 | 0.00085 | 0.108 |
| 31 | 0.00092 | 0.00087 | 0.087 |
| 33 | 0.00092 | 0.00084 | 0.133 |
| 35 | 0.00092 | 0.00087 | 0.087 |
| 36 | 0.00092 | 0.00088 | 0.096 |
| 41 | 0.00092 | 0.00087 | 0.091 |

**Appendix Table 3:** P-values associated with simulation analyses for α, β, and γ diversity. Significant p-values are shown with asterisks (* represents significance at 0.05; ** represents significance at 0.001). β_M_ signifies the multiplicative model, whereas β_A_ signifies the additive model. Simulated viral metacommunities were generated using three separate methods (columns), as described in the Appendix.

| Diversity effect | Sampling Without Replacement | Sampling With Replacement | Two-Step Sampling Process |
| --- | --- | --- | --- |
| $\alpha$ - sex | 0.103 | 0.102 | 0.091 |
| $\alpha$ - species | <0.001** | <0.001** | <0.001** |
| $\alpha$ - species*sex | <0.001** | <0.001** | <0.001** |
| $\beta_m$ - sex | 0.021* | 0.027* | 0.027* |
| $\beta_m$ - species | 0.576 | 0.430 | 0.595 |
| $\beta_a$ - sex | 0.881 | 0.884 | 0.878 |
| $\beta_a$ - species | 1.000 | 0.999 | 0.999 |
| $\gamma$ - sex | 0.874 | 0.870 | 0.864 |
| $\gamma$ - species | 1.000 | 1.000 | 1.000 |

**Appendix Figure 1:**
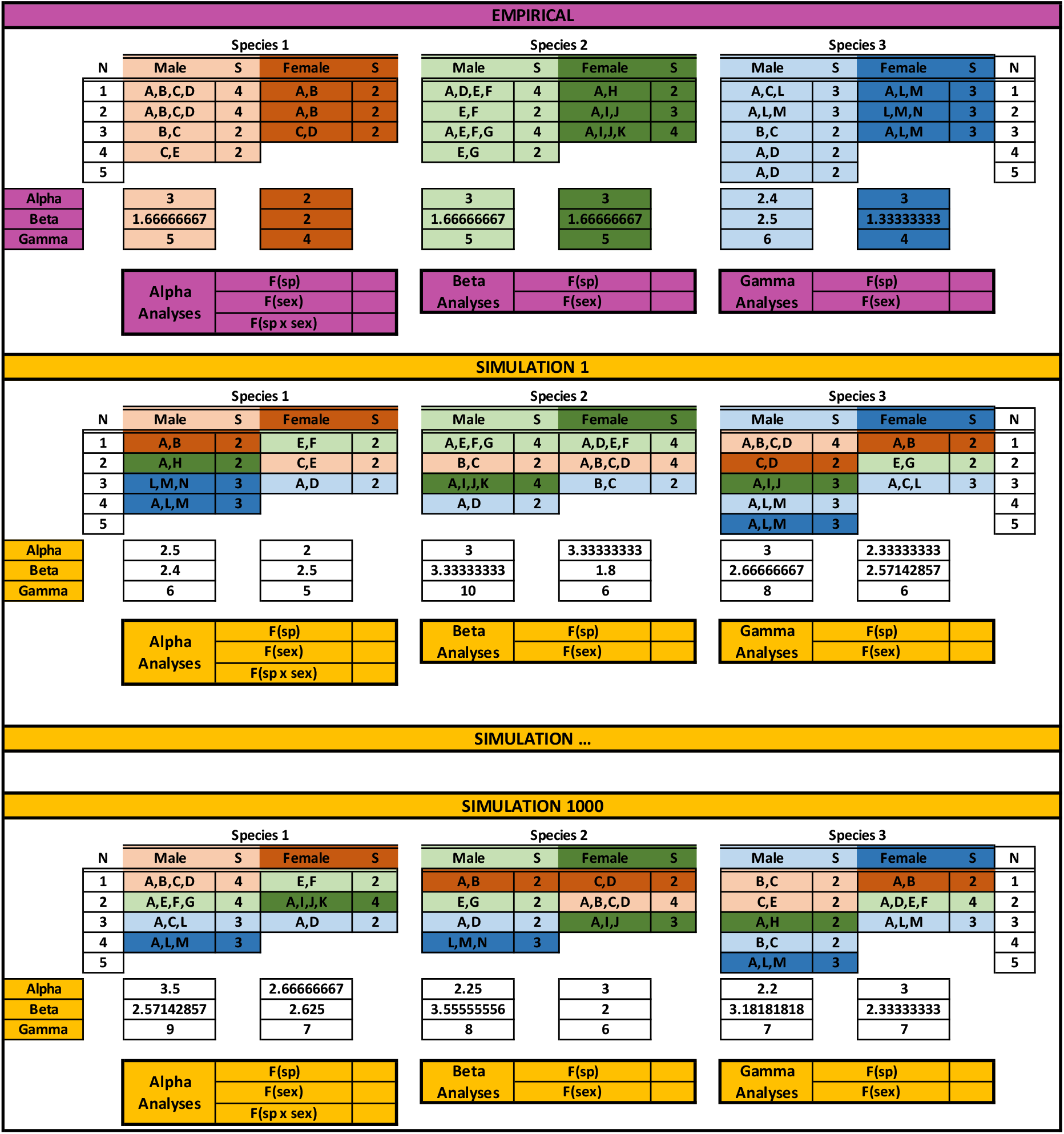
Simulation analyses for α, β, and γ diversity are shown in a simplified visualization. Example host species are represented by unique number and color combinations (Species 1 is orange, Species 2 is green, and Species 3 is blue), whereas example host sex is represented by shade of color (light for males versus dark for females). Example viral OTUs are represented by letters (A-M). For each simulation, number of hosts sampled per treatment group (N) remained equal to empirical numbers. Empirical patterns of OTU co-occurrence, and therefore OTU richness per host individual (S), also remained identical to that of the empirical data. First, for 900 iterations, a posterior OTU-host matrix prediction (purple) were used to calculate α, β, and γ diversity for each host treatment group, and appropriate F statistics were calculated. Then, for every simulation (1-1000, yellow), sampled infracommunities were randomly drawn from the pool and placed into a treatment group. F statistics were calculated for each level of analysis, and simulated F statistics were used to create null distributions. Empirical F statistics were then compared to the null distribution.

